# Structure of complement C3(H_2_O) revealed by quantitative cross-linking/mass spectrometry and modeling

**DOI:** 10.1101/056457

**Authors:** Zhuo A. Chen, Riccardo Pellarin, Lutz Fischer, Andrej Sali, Michael Nilges, Paul N. Barlow, Juri Rappsilber

## Abstract

The slow but spontaneous and ubiquitous formation of C3(H_2_O), the hydrolytic and conformationally rearranged product of C3, initiates antibody-independent activation of the complement system that is a key first line of antimicrobial defense. The structure of C3(H_2_O) has not been determined. Here we subjected C3(H_2_O) to quantitative cross-linking/mass spectrometry (QCLMS). This revealed details of the structural differences and similarities between C3(H_2_O) and C3, as well as between C3(H_2_O) and its pivotal proteolytic cleavage product, C3b, which shares functionally similarity with C3(H_2_O). Considered in combination with the crystal structures of C3 and C3b, the QCMLS data suggest that C3(H_2_O) generation is accompanied by the migration of the thioester-containing domain of C3 from one end of the molecule to the other. This creates a stable C3b-like platform able to bind the zymogen, factor B, or the regulator, factor H. Integration of available crystallographic and QCLMS data allowed the determination of a 3D model of the C3(H_2_O) domain architecture. The unique arrangement of domains thus observed in C3(H_2_O), which retains the anaphylatoxin domain (that is excised when C3 is enzymatically activated to C3b), can be used to rationalize observed differences between C3(H_2_O) and C3b in terms of complement activation and regulation.

## Introduction

The complement system performs immune surveillance, enabling rapid recognition and clearance of invading pathogens as well as apoptotic cells and particles threatening homeostasis (1). Multiple complement-activation pathways converge at the assembly of C3 convertases (2). These bimolecular proteolytic enzymes excise the anaphylatoxin domain (ANA, corresponding to C3a) from the complement component C3 (184 kDa) leaving its activated form, C3b (175 kDa) (Fig. 1A). C3b can covalently attach, *via* a nascently exposed and activated thioester, to any nearby surface (3, 4) whereupon it undergoes rapid amplification (2).

**Figure 1:**
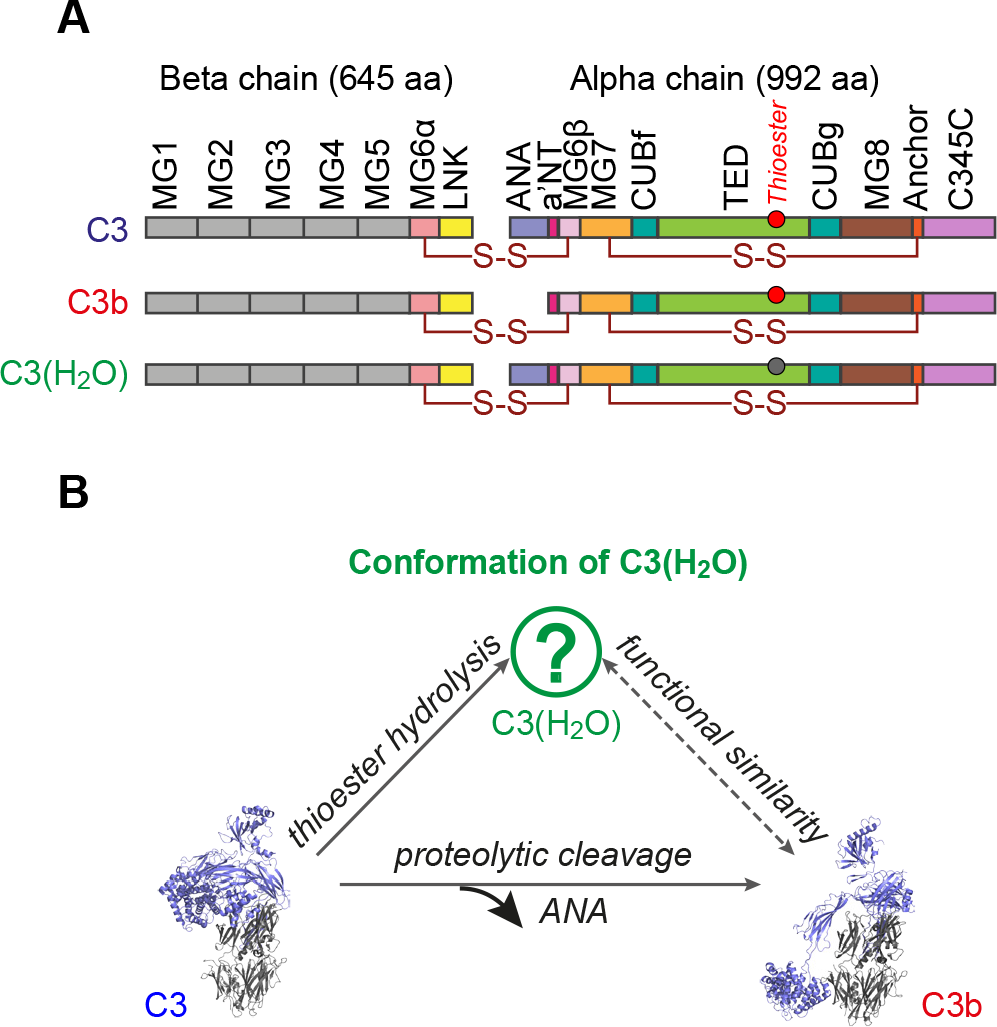
Complement protein C3(H_2_O) **(A)** Domain compositions of C3, C3b and C3(H_2_O). The thioester group in the TED is shown as a circle before (red) or after (gray) hydrolysis (25). **(B)** Relationship between native C3, C3(H_2_O) and C3b. The structure of C3(H_2_O) is unknown.

The complement system responds very swiftly to pathogens, independently of antibodies, due primarily to its “alternative pathway” of activation. This is initiated by spontaneous, although rare, conformational changes within C3 that are concerted with hydrolysis of its constitutively buried thioester linkage (5). Identical conformational changes accompany attack of the thioester by amines (6). The continuously and ubiquitously generated stable product, C3(H_2_O) (iC3 or C3N) does not bind to surfaces (as it no longer possesses a thioester group). Interestingly, C3(H_2_O) has been inferred to resemble C3b in many of its functional and structural features, despite its retention of the ANA (7–10) (Fig. 1A, B).

The mature C3 molecule consists of two polypeptide chains (residues 1-645 in the β-chain and residues 650-1641 in the α-chain). A metaphor of a puppeteer holding a puppet has been used to describe the crystal structure of C3 (11) (Supplemental Fig. S1 in Supplemental File). Macroglobulin domains (MGs) 1-6 and a “linking region” (LNK) adopt a key-ring like arrangement that forms the body of the puppeteer while MG7, MG8 and ANA form its shoulders, and a C345C domain equates to its head, joined to MG8 by an “anchor” region (the neck). A thioester-containing domain (TED) is the puppet, held at shoulder height by a CUB domain that forms the arm of the puppeteer. MGs 1-5, LNK and half of MG6 are contributed by the β-chain while the remaining domains are coming from the α-chain.

Comparing the crystal structures of C3 and C3b revealed significant domain rearrangements between them (11). Most dramatically, the CUB arm swings away from the shoulders towards the “feet” of the puppeteer (Supplemental Fig. 1S). As a result the TED (*i.e.* the puppet) rotates and is repositioned. This is accompanied by exposure and activation of the thioester group, allowing attachment of C3b to surface-borne nucleophiles. The crystal structure of C3(H_2_O) has not been reported. New binding sites for complement components and cell-surface receptors are created in both nascent C3b and C3(H_2_O) (7, 12–18). Both proteins bind factor B that is subsequently cleaved to Bb. Importantly, both the resultant C3bBb and C3(H_2_O)Bb complexes are C3 convertases, generating further molecules of C3b and thereby stoking a positive-feedback loop.

Because C3(H_2_O) (unlike C3b) is a spontaneously arising product of C3 domain rearrangements and thioester hydrolysis, C3(H_2_O)Bb (rather than C3bBb) is the initiating convertase of the alternative pathway of complement activation. Thus the constitutive presence of C3(H_2_O) ensures the alternative pathway can be activated quickly and indiscriminately allowing a rapid response to any cell not protected by the appropriate regulatory molecules such as factor H. Inappropriate regulation of complement activity is linked to many autoimmune, inflammatory and ischemia/reperfusion (I/R) injury-related diseases (19).

It has been shown that hydrolysis of the thioester in C3 alone does necessarily result in transition to active C3(H_2_O) (20). Despite use of diverse methodologies (7, 9–13, 21–27), the remodeling of domains that underlies spontaneous formation of C3(H_2_O), and therefore triggers complement, are poorly understood. Current structural models of C3(H_2_O) rely on epitope-mapping (21), hydrogen-deuterium exchange (27), other biophysical solution studies (9) and negative-staining EM images (25). These indicate a “C3b-like” structure but do not provide direct evidence regarding placements of the ANA and TED relative to specific domains within the shoulders and body of the C3(H_2_O) molecule. It has been proposed that the ANA domain acts as a safety catch in native C3. Removal of the ANA triggers the dramatic structural transition into C3b (24). More knowledge of the C3(H_2_O) structure is required to test if the safety catch role of ANA (presumably displaced in C3(H_2_O) rather than removed, as in C3b) and subsequent domain reconfigurations are general mechanisms, relevant both to the spontaneous but rare hydrolytic C3 to C3(H_2_O) transition, and to the proteolytic cleavage-dependent but rapid C3 to C3b transition.

Further understanding of this event depends on the ability to elucidate, in solution, the dynamic processes whereby the domains of a protein molecule are reorganized, following a triggering event, to form a new stable arrangement. Quantitative cross-linking/mass spectrometry (QCLMS) using isotope-labeled cross-linkers (Fig. 2A) has emerged as a new approach with which to elucidate the details of protein conformational changes (28–31). In this approach, chemical cross-linking captures proximities between amino acid residues and the residues involved are identified by mass spectrometry. Quantitative comparison of the crosslinking results obtained for two different conformations of a protein allows the details of the conformational change to be elucidated. We have developed a workflow for QCLMS analysis (32). In our benchmark study, we used QCLMS to accurately reveal differences and similarities between C3 and C3b in terms of the spatial arrangements of their domains (32). In another application, this technique successfully revealed conformational changes involved in maturation of the proteasome lid complex (33).

**Figure 2:**
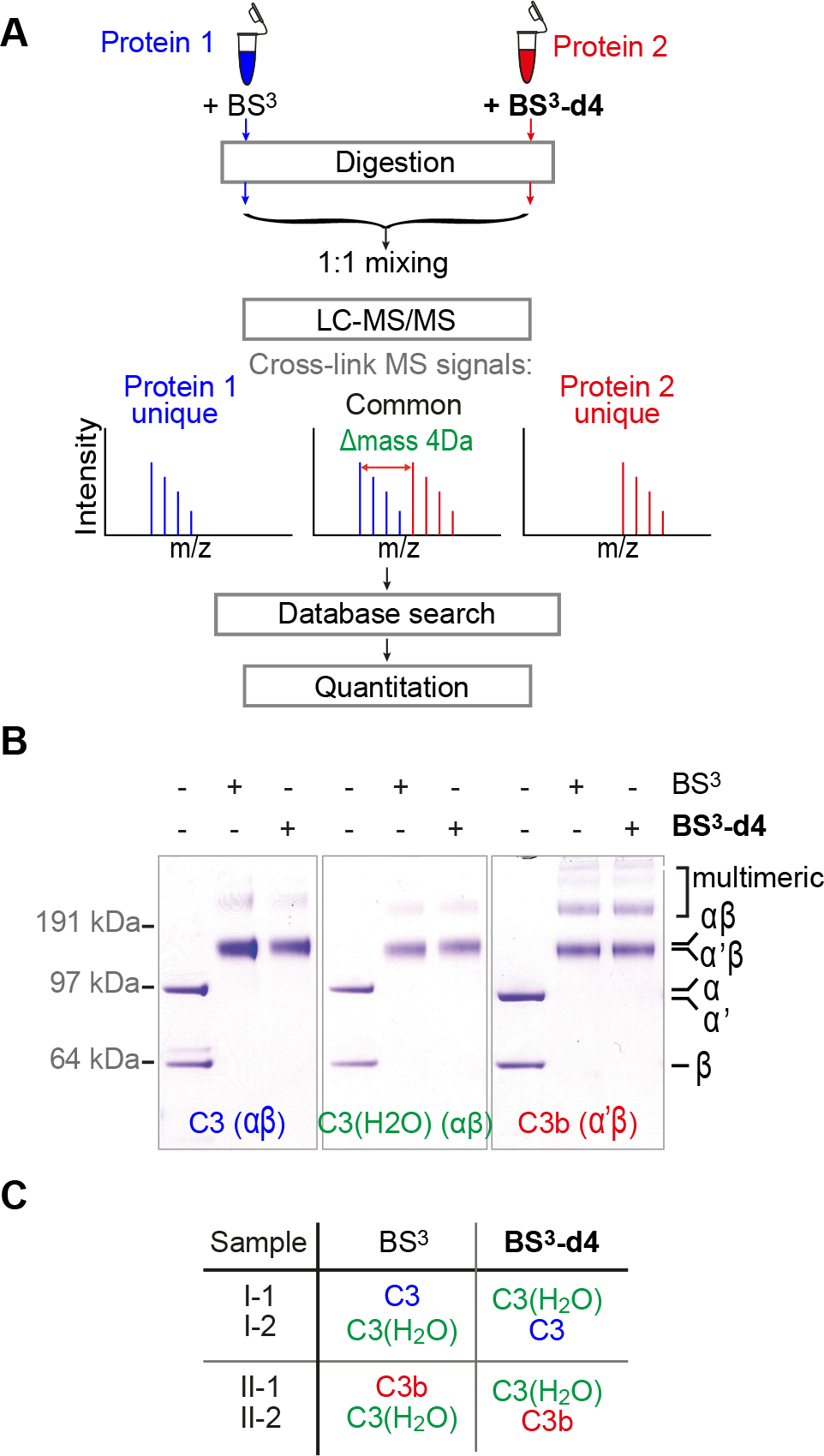
Quantitative CLMS analysis of C3, C3b and C3(H_2_O) in solution. **(A)** The strategy of QCLMS using differentially isotope-labelled cross-linkers for comparing protein conformations. **(B)** SDS-PAGE shows that BS^3^ (light cross-linker) and BS^3^-d4 (heavy cross-linker) cross-link C3, C3(H_2_O) and C3b with roughly equivalent overall efficiencies, and that broadly similar sets of cross-linked products were obtained. **(C)** Sample identifiers and color coding used in this study. For example sample I-1 consists of an equimolar mixture of BS^3^- cross-linked C3 (blue) and BS^3^-d4-cross-linked C3(H_2_O) (green). Pair-wise comparisons of C3(H_2_O) against C3 (I-1 and I-2), and C3(H_2_O) against C3b (red) (II-1 and II-2).

Here we apply our QCLMS workflow, and an integrative modeling approach, to interrogate the unknown arrangement of domains in, C3(H_2_O), a key component of the complement alternative pathway. We combined knowledge of the crystal structures of C3 and C3b with QCLMS datasets for C3(H_2_O), C3 and C3b. We thus generated structural models for the conformational transition of C3 to C3(H_2_O) that are consistent with other biophysical studies and with previously observed functional similarities and differences between these proteins.

## Experimental procedures

### Protein preparation for cross-linking

Plasma-derived human C3 and C3b were purchased from Complement Technology, Inc. USA (and stored at −80 °C). Native C3 was depleted of low amounts of contaminating C3(H_2_O) using cation-exchange chromatography (34). C3(H_2_O) (*i.e.* C3(N) in this case but identical to C3(H_2_O)) was prepared by incubating C3 at 37 °C with 200 mM methylamine (CH3NH2) at pH 8.3 for three hours. The C3(H_2_O) was then isolated from any other intermediates using cation-exchange chromatography (34). Chromatography in both cases was performed using a Mini S PC 3.2/3 column (GE Healthcare) at a flow-rate of 500 μl/min at 4 °C and a gradient from 0 to 325 mM NaCl. Immediately after purification, C3, C3(H_2_O) and C3b samples were exchanged, using 30-kDa molecular weight cutoff (MWCO) filters (Millipore), into cross-linking buffer (20 mM HEPES-KOH, pH 7.8, 20 mM NaCl and 5 mM MgCh) with a final concentration of 2 μM. C3, C3b and C3(H_2_O) samples were prepared in two separated batched and used for “experiment I” and “experiment II” respectively.

### Protein cross-linking

#### Experiment I

Fifty μg C3, C3b and C3(H_2_O) were each cross-linked in a volume of 100 uL with either bis[sulfosuccinimidyl] suberate (BS^3^) (Thermo Scientific) or its deuterated analogue bis[sulfosuccinimidyl] 2,2,7,7-suberate-d4 (BS^3^-d4) (Thermo Scientific), at 1:3 protein to crosslinker mass ratio, giving rise to six different protein-cross-linker combinations: C3+BS^3^, C3+BS^3^- d4, C3(H_2_O)+BS^3^, C3(H_2_O)+BS^3^-d4, C3b+BS^3^ and C3b+BS^3^-d4. After incubation (two hours) on ice, reactions were quenched with 10 μl 2.5 M ammonium bicarbonate for 45 minutes on ice. For the monitoring of cross-linking, aliquots containing 5 pmol of cross-linked protein from each of the above six reactions were subjected to SDS-PAGE using a NuPAGE 4-12% Bis-Tris gel (Life Technologies) and MOPS running buffer (Life Technologies). The protein bands were visualized using the Colloidal Blue Staining Kit (Life Technologies) (Fig. 2B). Cross-linking reactions were repeated for “experiment II” as described for “experiment I”.

### Sample preparation for mass spectrometric analysis

#### Experiment I

Each of C3, C3b and C3(H_2_O) consists of two polypeptide chains linked by a disulfide bond. The monomeric (two-polypeptide chains) product of cross-linked C3, C3b or C3(H_2_O) was isolated using SDS-PAGE (50 μg was loaded for each). Proteins were ingel reduced and alkylated, then digested using trypsin following a standard protocol (35). For quantitation, equimolar quantities of the tryptic products from the six cross-linked protein samples were mixed pair-wise in four combinations to allow the comparisons of C3(H_2_O) with C3b and with C3: C3+BS^3^ and C3(H_2_O)+BS^3^-d4 (sample I-1); C3+BS^3^-d4 and C3(H_2_O)+BS^3^ (sample I-2); C3b+BS^3^ and C3(H_2_O)+BS^3^-d4 (sample II-1); and finally C3b+BS^3^-d4 and C3(H_2_O)+BS^3^ (sample II-2) (Fig. 2C).

For each of the four samples, a 20 μg (40 μL) aliquot was fractionated using SCX-Stage-Tips (36) with a small variation of the protocol previously described for linear peptides (37). In short, peptide mixtures were first loaded on a SCX-Stage-Tip in loading buffer (0.5% v/v acetic acid, 20% v/v acetonitrile, 50 mM ammonium acetate). The retained peptides were eluted in two steps, with buffers containing 100 mM ammonium acetate and 500 mM ammonium acetate, into two fractions. These peptide fractions were desalted using C18-StageTips (38) prior to mass spectrometric analysis.

#### Experiment II

Preparation of four quantitation samples were repeated as described for “experiment I”. A 4-μg (10 μL) aliquot of each sample was desalted using C18-Stage-Tips for mass spectrometric analysis without pre-fractionation.

### Mass spectrometric analysis

#### Experiment I

SCX-Stage-Tip fractions were analyzed using a hybrid linear ion trap-Orbitrap mass spectrometer (LTQ-Orbitrap Velos, Thermo Fisher Scientific) applying a “high-high” acquisition strategy. Peptides were separated on an analytical column that was packed with C18 material (ReproSil-Pur C18-AQ 3 μm; Dr. Maisch GmbH, Ammerbuch-Entringen, Germany) in a spray emitter (75-μm inner diameter, 8-μm opening, 250-mm length; New Objectives) (39). Mobile phase A consisted of water and 0.5% v/v acetic acid. Mobile phase B consisted of acetonitrile and 0.5% v/v acetic acid. Peptides were loaded at a flow-rate of 0.6 μl/min and eluted at 0.3 μl/min using a linear gradient going from 3% mobile phase B to 35% mobile phase B over 130 minutes, followed by a linear increase from 35% to 80% mobile phase B in five minutes. The eluted peptides were directly introduced into the mass spectrometer. MS data were acquired in the data-dependent mode. For each acquisition cycle, the mass spectrum was recorded in the Orbitrap with a resolution of 100,000. The eight most intense ions with a precursor charge state 3+ or greater were fragmented in the linear ion trap by collision-induced disassociation (CID).The fragmentation spectra were then recorded in the Orbitrap at a resolution of 7,500. Dynamic exclusion was enabled with single repeat count and 60-second exclusion duration.

#### Experiment II

Non-fractionated peptide samples were analyzed using a hybrid quadrupole-Orbitrap mass spectrometer (Q Exactive, Thermo Fisher Scientific). Peptides were separated on a reversed-phase analytical column of the same type as described above. Mobile phase A consisted of water and 0.1% v/v formic acid. Mobile phase B consisted of 80% v/v acetonitrile and 0.1% v/v formic acid. Peptides were loaded at a flow rate of 0.5 μl/min and eluted at 0.2 μl/min. The separation gradient consisted of a linear increase from 2% mobile phase B to 40% mobile phase B in 169 minutes and a subsequent linear increase to 95% B over 11 minutes. Eluted peptides were directly sprayed into the Q Exactive mass spectrometer. MS data were acquired in the data-dependent mode. For each acquisition cycle, the MS spectrum was recorded in the Orbitrap at 70,000 resolution. The ten most intense ions in the MS spectrum, with a precursor charge state of 3+ or greater, were fragmented by Higher Energy Collision Induced Dissociation (HCD). The fragmentation spectra were thus recorded in the Orbitrap at 35,000 resolution. Dynamic exclusion was enabled, with single-repeat count and a 60-second exclusion duration.

### Identification of cross-linked peptides

The raw mass spectrometric data files were processed into peak lists using MaxQuant version 1.2.2.5 (40) with default parameters, except that “Top MS/MS Peaks per 100 Da” was set to 20. The peak lists were searched against C3 and decoy C3 sequences using Xi software (ERI, Edinburgh) for identification of cross-linked peptides. Search parameters were as follows: MS accuracy, 6 ppm; MS2 accuracy, 20 ppm; enzyme, trypsin; specificity, fully tryptic; allowed number of missed cleavages, four; cross-linker, BS^3^/BS^3^-d4; fixed modifications, carbamidomethylation on cysteine; variable modifications, oxidation on methionine; modifications by BS^3^/BS^3^-d4 that are hydrolyzed or amidated on the other end. The linkage specificity for BS^3^ was assumed to be for lysine, serine, threonine, tyrosine and protein N-termini. Identified candidates for cross-linked peptides were validated manually in Xi, after applying an estimated false discovery rate (FDR) of 3% for cross-linked peptides (Fischer and Rappsilber, submitted). We used 3% FDR as this was the best FDR that returned a reasonable number of decoys to provide a meaningful FDR. Only those cross-linked peptide pairs identified with fragment signals of both peptides in MS2 spectra were used to generate the list of identified cross-linked residue pairs and used for subsequent quantitation. Identification information of all quantified cross-linked peptides and the annotated best-matched MS2 spectra for these crosslinked peptides are provided in Supplemental Table S1 and Supplemental File S1.

### Quantitation of cross-link data using Pinpoint software

Quantitation was carried out in each pair-wise comparison. For each cross-linked peptide, the elution peak areas of light (BS^3^ cross-linked) and heavy (BS^3^-d4 cross-linked) signals were retrieved using Pinpoint (Thermo Fisher Scientific) (32, 33). The error tolerance for precursor m/z was set to 6 ppm. Signals were only accepted within a window of retention time (defined in spectral library) ±10 minutes. Manual inspection was carried out to ensure the correct isolation of elution peaks, and correct isotope peaks that were used for quantitation. “Match between runs” (41) was performed manually using Pinpoint software based on high m/z accuracy and reproducible chromatographic retention time for MS1 signals. Thus, signals of each identified cross-linked peptide were quantified in every quantitation samples. All transferred identification were verified based on their MS1 signal pattern (either shown as doublet signals or singlet signals with 4D mass shift between paired label-swapped replicas).

Differences between the yields of cross-linked peptide pairs were expressed in terms of “signal fold-changes” (i.e. by how many-fold the two signals differed). The signal fold-change of a cross-linked peptide pair was calculated as log2 (C3/C3(H_2_O)), or log2 (C3b/C3(H_2_O)). Within each quantitation sample, signal fold-changes of all observed cross-linked peptide pairs were first normalized to their median. This corrected systematic errors introduced by minor differences in mixing ratios during sample preparation. Then the signal fold-change for a residue pair was calculated as the median of all its supporting cross-linked peptides. Only those crosslinks that were consistently quantified in both paired replicas (i.e. with label-swapping) were accepted for subsequent structural analysis and the average of signal fold-changes of a residue pair from replicated analyses was calculated. When a cross-linked residue pair was quantified in both experiment I and experiment II, the average of signal fold-changes in two experiments was reported. All quantified cross-links are listed in Supplemental Table S2. Within each pair-wise comparison, the “Significance A” test from the standard proteomics data analysis tool Perseus (version 1.4.1.2) (40) was carried out based on fold-change values to determine cross-links that are significantly enriched in either conformations. The following parameters were used for the test: “Side”: both; “Use for truncation”: P value; “Threshold value”: 0.05.

### Visualizing cross-linking data in crystal structures

PyMol (version 1.2b5) (42) was used to visualize cross-linking data. Cross-links were displayed in the crystal structures of C3 (PDB|2A73) and C3b (PDB|2I07) as dashed lines between the C-α atoms of linked residues. In the case of a residue missing from the crystal structures, the nearest residue in the sequence was used for display purposes. The distance of a cross-linked residue pair in the crystal structures was measured between the C-α atoms. The theoretical cross-linking limit was calculated as the sum of side-chain lengths of cross-linked residues plus the spacer length (11.4 Å) of the cross-linker. An additional 2 Å were added for each residue to allow for residue displacement in the crystal structures. The following side-chain lengths were used for the calculation: 6.0 Å for lysine, 2.4 Å for threonine, 2.4 Å for serine and 6.5 Å for tyrosine. For example, for a lysine-lysine cross-link, this limit is 27.4 Å.

### Determining the structures of C3(H_2_O), C3, C3b with Integrative Modeling Platform (IMP)

Our integrative approach for determining the structure of the three macromolecules proceeded as follows (43–48): (1) gathering of data, (2) representation of subunits and translation of the data into spatial restraints, (3) configurational sampling to produce an ensemble of models that optimally satisfies the restraints, and (4) analysis and assessment of the ensemble. The modeling protocol was scripted using the Python Modeling Interface (PMI), a library for modeling macromolecular complexes based on the open-source IMP package (46), release 2.5.0. The protein domains were represented as rigid or flexible, based on known protein structures. The cross-linking data (sets of 82, 75, and 85 cross-links for C3, C3b, and C3(H_2_O), respectively. Supplemental Table S3) were encoded into a Bayesian scoring function that restrained the distances spanned by the cross-linked residues (44, 45). We also included the disulfide bond between residues 851 and 1491. Models of C3, C3b, and C3(H_2_O) were computed separately. The 200 best scoring models (i.e., solutions) for each protein were clustered to yield the localization density maps (further described below). The average precision of the solutions for C3(H_2_O) (average r.m.s. (root-mean-square) deviation with respect to the cluster center when superposed on the MG1-6_α’NT domain) was 15 Å (Fig. 6E, green bars). The precision of the domains varies because the intra-domain cross-links are not uniformly distributed. The C-alpha r.m.s. deviation of the solutions with respect to the crystallographic structures (PDB 2I07 and 2A73) was used to estimate the accuracy of the C3 and C3b models (Fig. 6E, blue and red bars). The C3 and C3b crystallographic structures were reproduced with an accuracy of approximately 11 Å.

### Representation of domains

The domains of the complement protein C3 were represented by beads arranged into either a rigid body or a flexible string on the basis of the available crystallographic structures (PDB 2I07 for C3b and PDB 2A73 for C3 and C3(H_2_O)) (Fig. 6A). The beads representing a structured region were kept rigid with respect to one another during configurational sampling (i.e., rigid bodies). Segments without a crystallographic structure or the linkers between the rigid domains were represented by a flexible string of beads, where each bead corresponded to a single residue.

### Bayesian scoring function

The Bayesian approach estimates the probability of a model, given information available about the system, including both prior knowledge and newly acquired experimental data. The approach is extensively described elsewhere (44, 45, 49, 50). Briefly, using Bayes’ theorem, we
estimate the posterior probability *p*(*M*|*D*,*I*), given data *D* and prior knowledge *I*, as *p*(*M*|*D*, *I*) ∝ *p*(*D*|*M*, *I*)*p*(*M*, *I*), where the likelihood function p(D\M,I) is the probability of observing data *D*, given *I* and *M*, and the prior is the probability of model *M*, given *I*. To define the likelihood function, one needs a forward model that predicts the data point (i.e., the presence of a crosslink between two given residues) given any model *M* and a noise model that specifies the distribution of the deviation between the observed and predicted data points.

To account for the presence of noisy cross-links, we parameterized the likelihood with a set of variables {Ψ} defined as the uncertainties of observing the cross-links in a given model (44, 45). YC is the average uncertainty for cross-links that were consistently identified in all cross-linking experiments, ΨI is the average uncertainty for cross-links that were identified only once. For instance, cross-link 1049^TED^-1409^MG8^ was identified for C3 in both C3(H_2_O)/C3 and C3b/C3 quantification experiments, therefore its uncertainty was estimated by ΨC. Conversely, the cross-link 203^MG2^-1049^TED^ was identified for C3 in C3b/C3 but not in C3(H_2_O)/C3 and its uncertainty was identified by ΨI.

The prior terms comprised the excluded volume and the sequence connectivity, which are described elsewhere (44, 45). Moreover, the disulfide bond is implemented as a harmonic restraint on the distance between the two cross-linked residues.

### Sampling model configurations

Structural models were obtained by Replica Exchange Gibbs sampling, based on Metropolis Monte Carlo sampling (50). This sampling was used to generate configurations of the system as well as values for the uncertainty parameters. The Monte Carlo moves included random translation and rotation of rigid bodies (4 Å and 0.03 rad, maximum, respectively), random translation of individual beads in the flexible segments (5 Å maximum), and a Gaussian perturbation of the uncertainty parameters. The sampling was run on 32 replicas, with temperatures ranging between 1.0 and 2.5. Two independent sampling calculations were run for each system, each one starting with a random initial configuration, for a total of 200,000 models per system. We divided this set of models into two ensembles of the same size to confirm sampling convergence (data not shown).

### Analysis of the model ensemble

For each ensemble, the solutions were grouped by *k*-means clustering on the basis of the r.m.s. deviation of the domains after the superposition of the MG1-6_α_’NT domain (Supplementary Fig. S3 in Supplemental File). For C3 and C3b, the models cluster into a single configuration. For C3(H_2_O), the models cluster into two configurations (A and B). We chose the cluster with that best satisfied the cross-linking data. The precision of a cluster was calculated as the average r.m.s. deviation with respect to the cluster center (i.e., the solution with the lowest r.m.s. deviation with respect to the others). The solutions of a cluster, superposed on the MG1-6_α_’NT domain, were converted into the probability of a volume element being occupied by a given domain (i.e. the localization density, Fig. 6B, C, D) (51).

### Accession codes

IMP modelling scripts and models are available at https://mycore.core-cloud.net/public.php?service=files&t=09219adca568ac091ca148410a4ba6da

The mass spectrometry proteomics data have been deposited to the ProteomeXchange Consortium (52) (http://proteomecentral.proteomexchange.org) via the PRIDE partner repository with the dataset identifier PXD003486.

## Results

### C3(H_2_O) *versus* C3 comparison confirms structural rearrangements in C3(H_2_O)

To interrogate the unknown arrangement of domains in C3(H_2_O), we first compared QCLMS data for C3(H_2_O) and C3 (for which a crystal structure has been determined). Using our tried and tested workflow (Fig. 2A) (32, 33), a total of 94 cross-linked pairs of amino acid residues (“cross-links”) were quantified for these two proteins (Fig. 3A). Ten cross-links were identified uniquely in C3 and 22 cross-links were found only in C3(H_2_O). Of 62 cross-links that were observed in both C3 and C3(H_2_O), one was significantly enriched in C3 compared to C3(H_2_O) and four were significantly enriched in C3(H_2_O) compared to C3 (determined using the “Significance A” test from Perseus version 1.4.2.1 with *p*<0.05) (40). The observation of 37 (39% of the total) cross-links differing in occurrence or in enrichment between C3(H_2_O) and C3 reflects significant structural differences between these proteins. Yet these proteins must also have structural features in common, given that 57 out of 94 cross-links showed no significant differences in yields between the two proteins.

**Figure 3:**
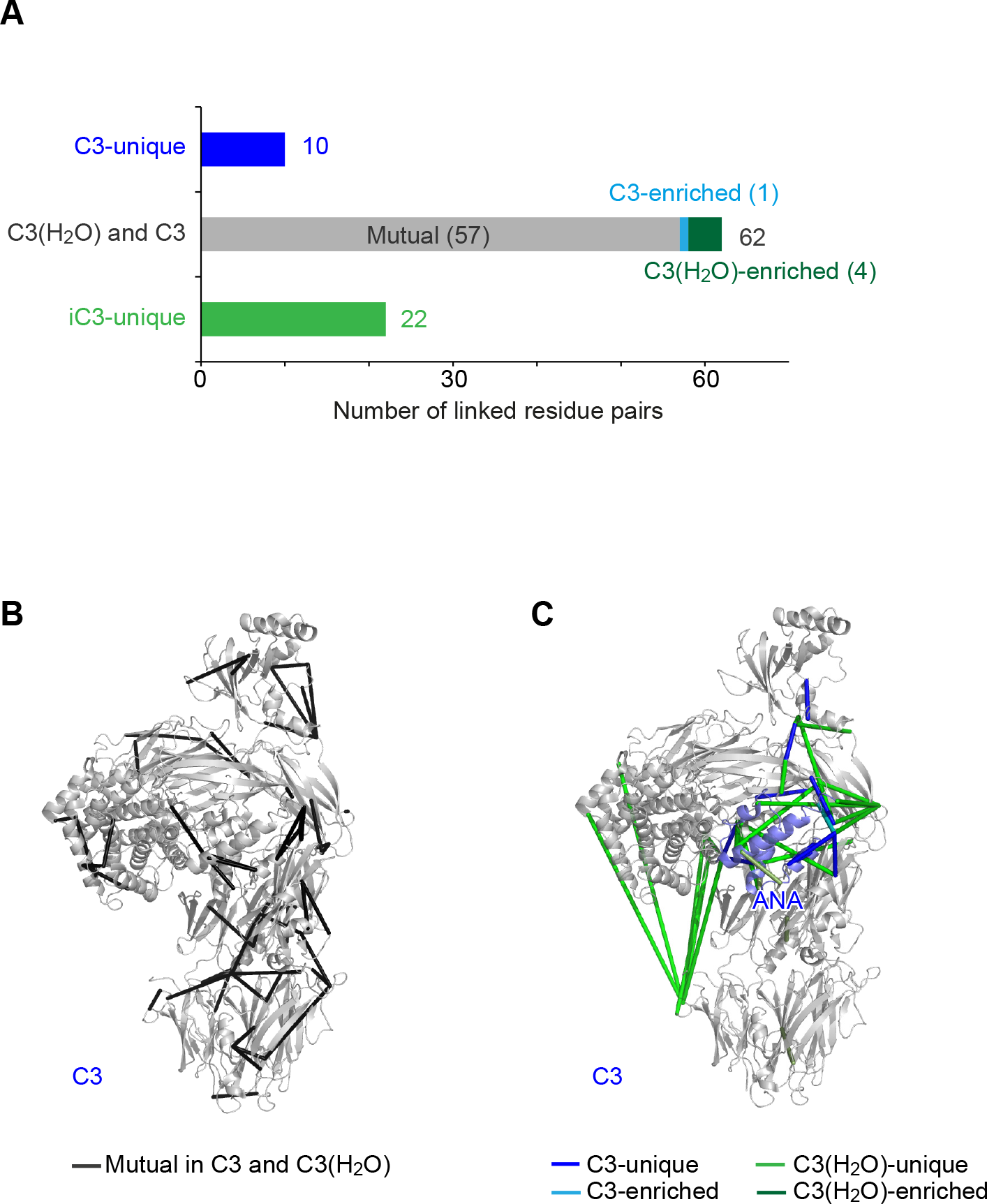
Comparison of cross-links in C3(H_2_O) and C3. The 94 cross-links quantified from the C3(H_2_O) *versus* C3 comparison fall into five groups: C3-unique (10), C3-enriched (1), C3(H_2_O)-C3 mutual (57), C3(H_2_O)-enriched (4) and C3(H_2_O)- unique (22). **(B)** The 57 C3(H_2_O)-C3 mutual cross-links are displayed (as dark grey rods) on the crystal structure of C3 (PDB|2A73), reflecting the presence of similar structural features. **(C)** Ten C3-unique cross-links (blue rods), one C3-enriched cross-link (cyan rod), four C3(H_2_O)-enriched cross-links (dark green rods) and 22 C3(H_2_O)-unique cross-links (green rods) are displayed on the crystal structure of C3, suggesting the locations of structural differences between C3(H_2_O) and C3. ANA is colored in light blue.

The 57 cross-links that are preserved upon the formation of C3(H_2_O) from C3 (referred to hereafter as “C3(H_2_O)-C3 mutual” cross-links) were inspected to identify structural features shared between the activated (C3(H_2_O)) and non-activated (C3) versions of this protein. Altogether 29 of these 57 cross-links, covering ten domains, connect residues within the same domains (Fig. 3B). It seems likely that these domains retain their structures following transition of C3 to C3(H_2_O). Furthermore, out of the total of 16 cross-links between domains within the β-chain, 15 are C3(H_2_O)-C3 mutual, suggesting that the respective β-chains share highly similar domain architectures. By contrast, the ten C3(H_2_O)-C3 mutual cross-links between domains in the α-chain account for only about 35% of the inter-domain cross-links identified in the α-chain. Thus, following transition to C3(H_2_O), the α-chain undergoes more conformational adjustments than the β-chain. Only three of twelve cross-links between the α-chain and the β-chain (the MG6p segment was treated as part of the β-chain for the purposes of this analysis) are C3(H_2_O)-C3 mutual. In the crystal structure of C3, all three are located at the interface between the α-chain and the β-chain, linking MG3 and MG7 (Fig. 3B).

From the above, it is clear that non-(C3-C3(H_2_O)) mutual cross-links, indicating structural differences, occur predominantly in the α-chain. They are concentrated in two distinct regions. The first group of C3-specific or C3(H_2_O)-specific cross-links occur in the vicinity of ANA and its neighboring domains in the “shoulder” region of the molecule (Fig. 3C). Inspection of crosslinking networks implied close proximities between these domains in both C3 and C3(H_2_O). Yet nine C3-unique (and enriched) cross-links and 17 C3(H_2_O)-unique (and enriched) cross-links were found within this region (Fig. 3C). This observation suggests that these domains undergo significant rearrangements while remaining intimately associated during the C3-C3(H_2_O) transition. Of these 26 non-mutual cross-links, five involve pairs of residues within the ANA domain and one involves a pair of residues within the MG8 domain. This observation implies that transition-associated conformational changes within these two domains accompany the more general domain rearrangements in the region. The second group of C3-specific or C3(H_2_O)-specific cross-links (Fig. 3C) revealed a drastic repositioning of TED relative to the main body of the molecule. In C3, TED was exclusively cross-linked to MG8 and MG3, at the “shoulder” of the molecule, while in C3(H_2_O) six unique cross-links suggest a newly established proximity between TED and MG1 at the “foot” of the molecule.

In summary, simple inspection of the QCLMS data reveals radical structural rearrangements in the α-chain of C3(H_2_O) compared to C3. The TED domain in particular was relocated. In addition, rearrangements at the “shoulder” region resulted in significant although less dramatic shifts in relative position of its component domains. In contrast, the conformation of the β-chain is largely preserved between C3 and C3(H_2_O). Such a structural transition is very similar to what was observed when comparing the crystal structure of C3 (24) with the crystal structure of its activated fragment form, C3b (11). Structural similarities between C3(H_2_O) and C3b were therefore investigated based on QCLMS data for both proteins.

### C3(H_2_O) *versus* C3b comparison confirms similar domain architectures

In total, 92 cross-links were quantified in the C3(H_2_O) *versus* C3b comparison (Fig. 4A). The detection of 57 C3(H_2_O)-C3b mutual cross-links, distributed throughout the structure of C3b, proves that these two activated derivatives of C3 have similar structures (Fig. 4B). As with the C3 *versus* C3(H_2_O) comparison, the arrangements of β-chain domains appear to be nearly identical, but C3(H_2_O) and C3b also share extensive structural features in their α-chains, and in the interfaces between their α-chains and β-chains.

**Figure 4:**
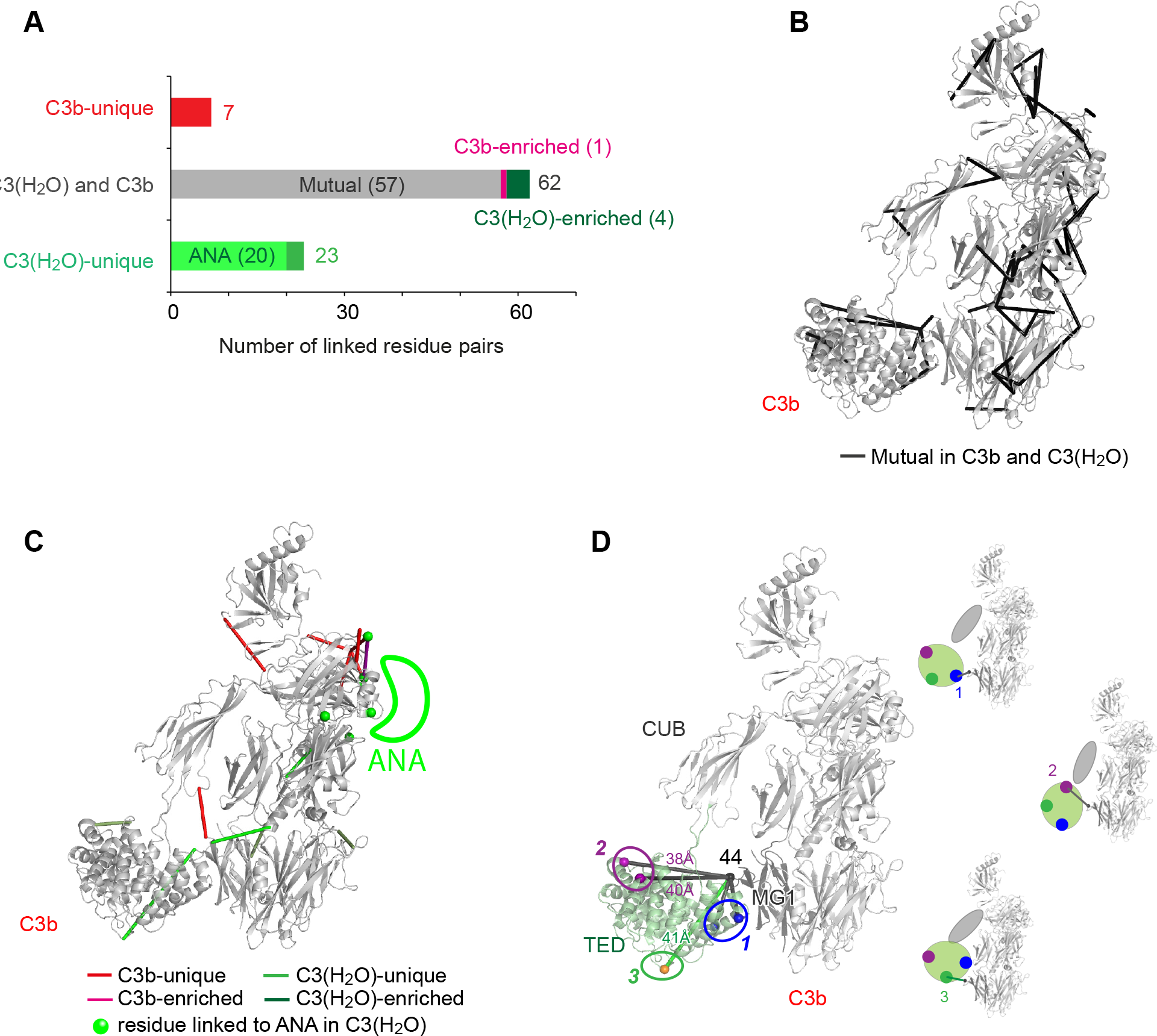
Comparison of cross-links in C3(H_2_O) and C3b. **(A)** The 92 cross-links quantified from the C3(H_2_O) *versus* C3b comparison fall into five groups: C3(H_2_O)-unique (23), C3(H_2_O)-enriched (4), C3(H_2_O)-C3b mutual (57), C3b-enriched (1) and C3b-unique (7). **(B)** The 57 C3(H_2_O)-C3b mutual cross-links are displayed (as dark grey rods) on the crystal structure of C3b (PDB|2I07), reflecting similar structural features. **(C)** Seven C3b-unique cross-links (red rods), one C3b-enriched cross-link (magenta rod), four C3(H_2_O)-enriched cross-links (dark green rods) and 16 C3(H_2_O)-unique cross-links (green rods) are displayed on the crystal structure of C3b, Note that C3(H_2_O) residues that are cross-linked to ANA (absent from C3b) are shown as green spheres. The likely position of the C3(H_2_O) ANA is indicated but the seven cross-links within ANA are not shown. These C3b or C3(H_2_O)-unique cross-links suggest the locations of structural differences between C3(H_2_O) and C3b. **(D)** Five cross-links from 44^MG1^ and TED (visualized in the C3b crystal structure PDB|2I07) indicate that three distinct sites on TED (marked as 1, 2, and 3) can be proximal with MG1, reflecting three different orientations of TED relative to the structural core (shown schematically in the three structures drawn at a smaller scale). Cross-links from MG1 to site 1 and 2 are mutual to C3b and C3(H_2_O), while the cross-link to site 3 is unique to C3(H_2_O). In the crystal structure of C3b, Cα-Cα distances between 44^MG1^ and cross-linked residues at site 2 (1181^TED^ and 1195^TED^) and site 3 (1049^TED^) violate the BS3 cross-link limit (27.4 Å). These observations suggest mobility of CUB and TED with respect to MG1-6 in both C3b and C3(H_2_O).

Of the total of 21 C3b-C3(H_2_O) mutual cross-links connecting α-chain residues to β-chain residues, eleven are also present in C3, while ten are not. Of these ten C3(H_2_O)-C3b mutual, but non-C3, cross-links, five connect TED and MG1. This supports suggestions that the TED domain in C3(H_2_O) migrates along the length of the molecule, as it does in C3b. This new domain arrangement likely creates a key functional interface, common to C3b and C3(H_2_O), for binding of proenzyme factor B (Fig. 5I) or regulator factor H (8, 10, 11, 53).

**Figure 5:**
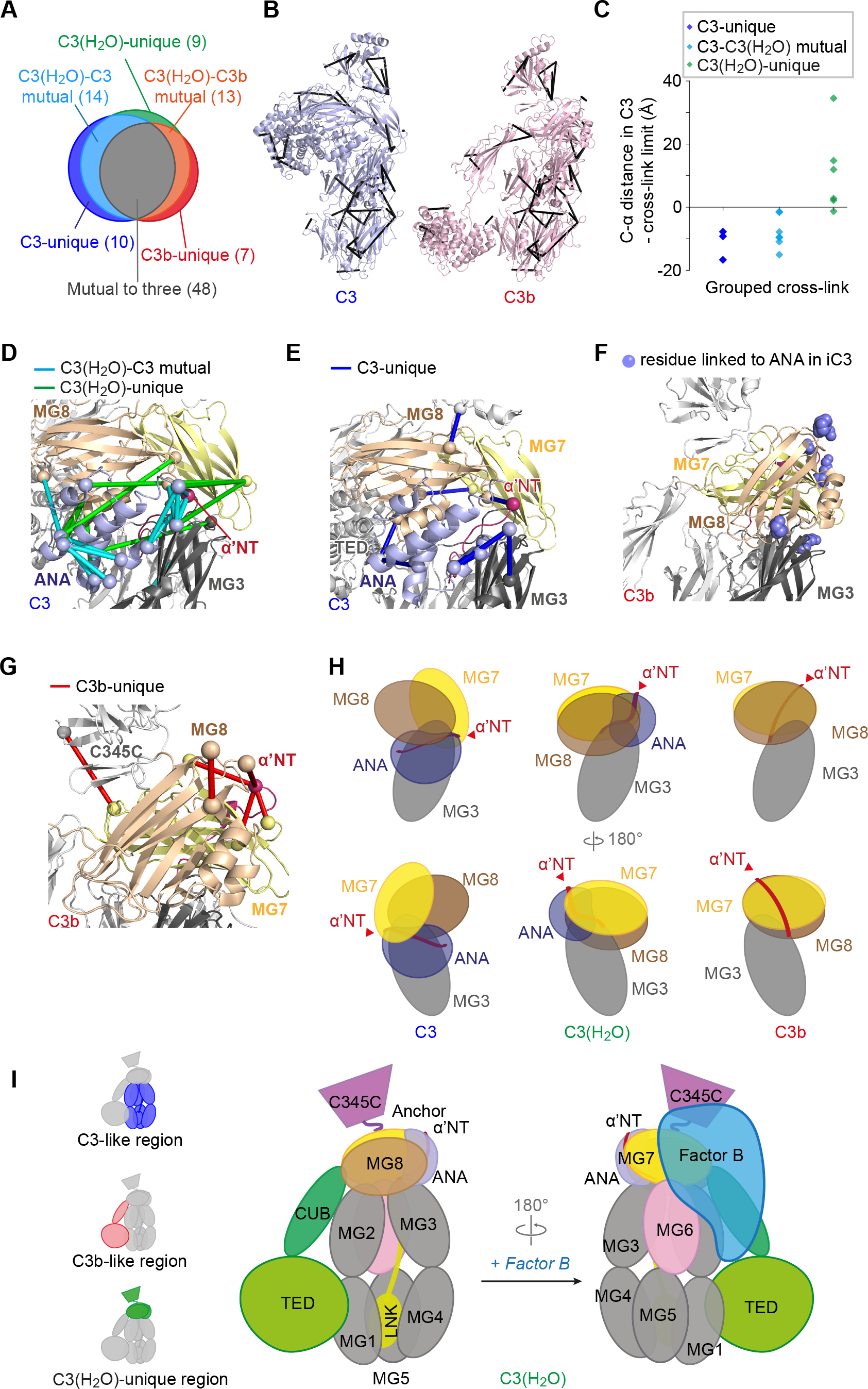
Comparison of cross-links in C3(H_2_O) and C3. **(A)** Combining pair-wise comparisons of C3(H_2_O) *versus* C3 and C3(H_2_O) *versus* C3b resulted in 101 quantified cross-links, falling into six categories as summarized in the Venn diagram. **(B)** The 48 cross-links that were observed in C3, C3b and C3(H_2_O) are drawn as black rods in the crystal structures of C3 (PDB|2A73) and C3b (PDB|2I07). These cross-links reflect structural features that are preserved in all three proteins. **(C)** Cα-Cα distances are shown for the three C3-unique, six C3(H_2_O)-unique and seven C3(H_2_O)-C3 mutual (but missing from C3b) crosslinks, as measured in the C3 structure (PDB|2A73). These cross-links all involve residues of ANA. The theoretically maximum cross-linkable distance was subtracted, and the data plotted, showing that most of the C3(H_2_O)-unique cross-links are incompatible with the relative position of ANA as seen in C3. **(D)** In the “shoulder” region of the C3 structure are shown 14 C3(H_2_O)- C3 mutual cross-links (cyan rods) and nine C3(H_2_O)-unique cross-links (green rods) (C-α atoms of cross-linked residues shown as spheres). The domains in this region of C3(H_2_O) must be rearranged relative to their positions in C3. **(E)** In the “shoulder” region of C3 are shown ten C3-unique cross-links (blue rods), indicating structural features of C3 that are likely absent in C3(H_2_O). **(F)** Cross-linking partners of ANA residues in C3(H_2_O) are highlighted (blue spheres) in the C3b crystal structure (PDB|2I07). This suggests a likely location of ANA relative to the rest of the C3(H_2_O) structure (cf. Fig. 3C). **(G)** Detailed view of six C3b-unique cross-links (red) within the “shoulder” region (four from Ser727, the α’ chain N-terminus in C3b) drawn on the C3b structure (PDB|2I07). These cross-links suggest structural features of C3b that are likely to be missing from C3(H_2_O). **(H)** A model (center) of a unique arrangement, in C3(H_2_O), for ANA, the neighboring MG7 and MG8 domains, and the region of C3(H_2_O) corresponding to α’-NT of C3b. The arrangements of these domains in C3 and C3b structures are shown for reference. **(I)** A schematic to represent the domain architecture of C3(H_2_O), inferred from the quantified crosslinking data and the crystal structures of C3 and C3b, with the putative binding site for factor B indicated. In the inset, the domains that are C3-like (blue) or C3b-like (red) in arrangement, or display features unique to C3(H_2_O) (green), are highlighted. This inferred structure can be compared with the integrative model in Fig. 6.(A)

**Figure 6.**
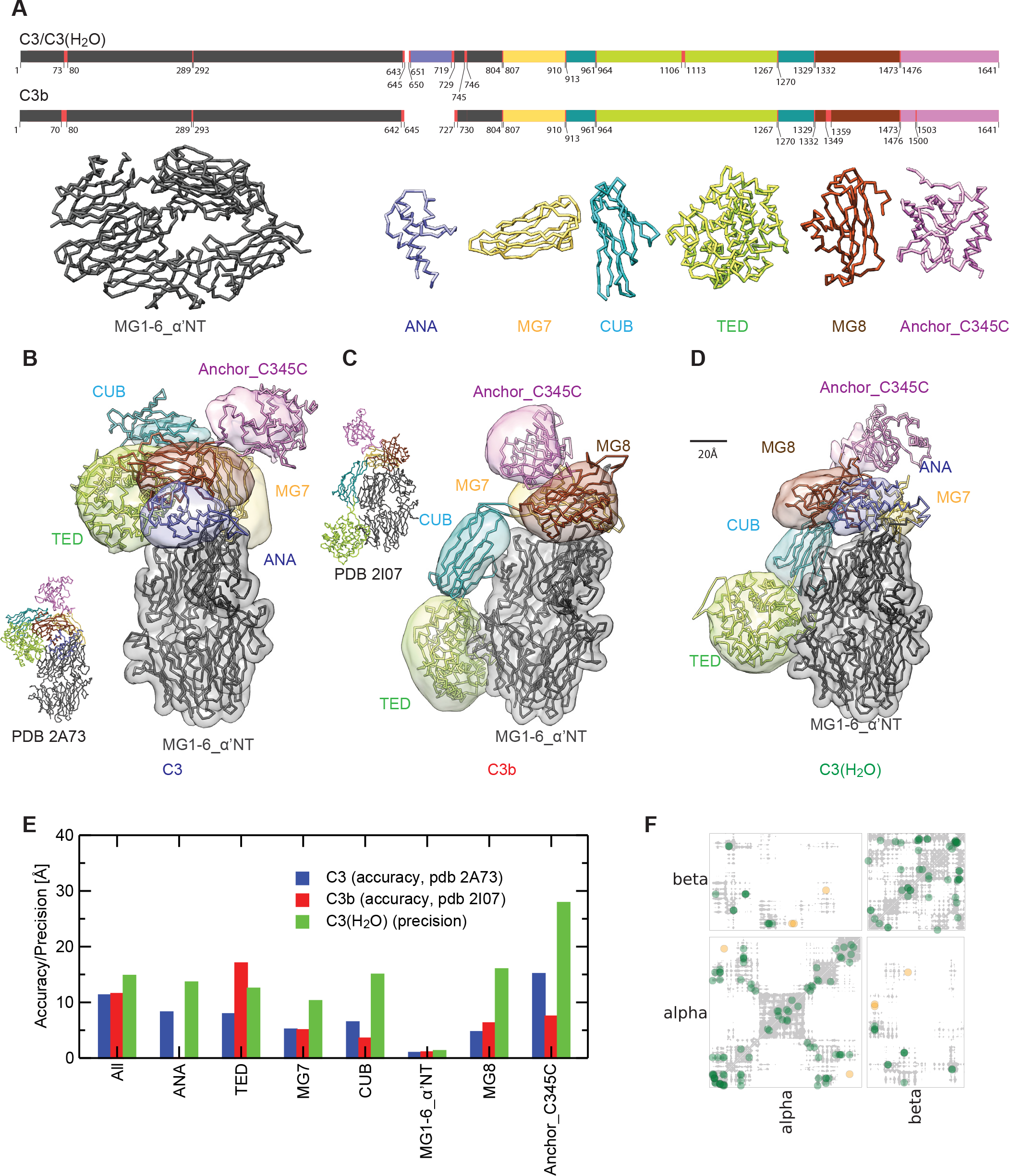
An integrative model for C3(H_2_O) **(A)** The positions and structures of domains treated as rigid bodies in our modeling of C3, C3b and C3(H_2_O) structures. Horizontal bars are color-coded to indicate the positions of domains/rigid bodies within the primary sequences. The red insertions were treated as flexible strings of beads that allow mobility between the rigid bodies. (**B-D**) The models of C3, C3b, and C3(H_2_O) (obtained as described in Experimental Procedures) are displayed using the localization densities of their domains (see (**A**)), shown as transparent surfaces. Backbone traces of models are also shown. The models of C3 and C3b compare well with the corresponding crystallographic structures (PDB|2A73 and PDB|2I07) that are shown alongside at a reduced scale. (**E**) The accuracy and the precision of C3, C3b, and C3(H_2_O) domains. The accuracy is the average root mean squared distance (r.m.s.d) (Cα atoms) of the cluster of solutions with respect to the corresponding crystallographic structure (PDB|2A73 for C3 or PDB|2I07 for C3b). The precision is the average r.m.s.d (Cα atoms) of the cluster of solutions with respect to the cluster center (defined in Experimental Procedures). Both accuracy and precision are computed after alignment of solutions on the rigid body MG1-6_α_’NT. **(F)** Contact map for cluster B of solutions of C3(H_2_O) model (grey shades) overlaid with the cross-links map (colored circles). Green and orange circles are satisfied and violated cross-links, respectively (a satisfied cross-link is defined when the Cα-Cα distance is below 35 Å (44)).

### The arrangement of domains in the “shoulder” region of C3(H_2_O)

In addition to similarities, there are also differences between the arrangements of domains in C3(H_2_O) and C3b as evidenced by our identification of 35 non-mutual cross-links. Not surprisingly, 20 of these involve the ANA domain, which is absent in C3b. Thirteen of these 20 cross-links are formed between ANA and the other four shoulder domains, α’-NT (N-terminal segment of the α’-chain in C3b formed by ANA cleavage), MG3, MG7 and MG8. These crosslinks provide unique information on the location of ANA in C3(H_2_O) and how this differs from its position in C3 (Fig. 4C). Of particular interest are the 15 C3b/C3(H_2_O)b non-mutual cross-links that are not attributable to ANA. Eight of these are unique to, or enriched in, C3b, while seven are unique to, or enriched in, C3(H_2_O). Four of the C3b-unique cross-links involve residue S727. This residue corresponds to the new N-terminus of the α’-chain in C3b formed by the cleavage of ANA. Five cross-links (three C3(H_2_O)-unique cross-links to the ANA domain plus two C3(H_2_O)-C3b mutual cross-links) involving S727 were observed in C3(H_2_O). Further information on the position of the new N-terminus emerged from the three-way comparison of C3(H_2_O), C3 and C3b cross-links described below.

### The positioning of TED in C3(H_2_O) versus C3b

Of note, two of the five C3(H_2_O)/C3b-mutual, MG1-TED cross-links correspond to residue-pairs that are far apart in the crystal structure of C3b (PDB|2I07); 38.3 and 40.4 Å, respectively, compared to a theoretical cross-linking limit of 27.4 Å (Experimental procedures). This could be explained if the TED domains of C3(H_2_O) and C3b were both mobile with respect to MG1 in solution, as also suggested by previous studies (25, 54, 55). Our data additionally suggest differences between C3(H_2_O) and C3b with respect to the organization of the CUB and TED domains relative to MG1. A cross-link 75^MG1^-1293^CUB^ was detected in C3b but was absent inC3(H_2_O). This agree with previous SAXS-based observation, which suggested that TED and CUB are positioned, on average, further away from the MG domains in C3(H_2_O), compared to what is seen in the various crystal structures of C3b (9). In addition we observed a C3(H_2_O)- unique cross-link 44^MG1^-1049^TED^. To enable cross-linking between this pair of residues, a CUB-TED position in respect to MG1 is required, which was not captured by cross-linking in C3b (Fig. 4D). Thus while TED is mobile, relative to MGs 1-6, in both C3(H_2_O) and C3b, it appears to be more mobile in C3(H_2_O) (9).

### Unique domain architecture of C3(H_2_O) based on a comparison of all three proteins

We next combined quantitation results from both the C3(H_2_O) *versus* C3 comparison and the C3(H_2_O) *versus* C3b comparison (Supplemental Table S2). In total 101 cross-links were quantified (Fig. 5A) in this comparison including nine that were unique to C3(H_2_O) and 48 that were mutual to all three proteins (Fig. 5B). None of the cross-links fell into the theoretical category of being mutual to C3 and C3b, but absent in C3(H_2_O). This analysis provided valuable additional insights into the architecture of C3(H_2_O).

The three-way comparison of cross-links reinforces the aforementioned inferences from the C3(H_2_O)-C3 comparison regarding ANA. Seven C3(H_2_O)-C3 mutual cross-links within this domain suggest a broadly C3-like ANA structure in C3(H_2_O) (Fig. 5D). But three cross-links within the N-terminal α-helix of ANA are unique to C3 (Fig. 5E). This, together with data between cross-links from ANA to MG3 and MG7 (discussed further below), implies that a conformational rearrangement accompanies C3(H_2_O) formation that involves both relocation of the ANA domain and a structural change within the N terminus of the ANA domain.

The three-way comparison also supplements the previously described two-way comparisons by suggesting that C3(H_2_O) formation from C3, like C3b formation from C3, involves a movement of MG8 towards MG3 (Fig. 5C, D, H). Specifically, a cross-link between MG3 and MG8 (267^MG3^-1409^MG8^) that had similar yields in both C3b and C3(H_2_O), was not detected in C3. In the C3 crystal structure, the Cα atoms of residues 267 and 1409 are 35 Å apart and therefore lie beyond the theoretical cross-linking limit for BS^3^ (27.4 Å). In C3b, these residues are 17.3 Å apart.

In the C3-to-C3b transition the approach of these residues is facilitated by proteolytic removal of the ANA domain. In contrast, for the case of the C3-to-C3(H_2_O) transition, the ANA domain remains attached and therefore this movement of MG8 towards MG3 would require it to be displaced. Indeed, three cross-links between ANA and MG7 occurred exclusively in C3(H_2_O), supporting a movement of the ANA domain towards MG7 in C3(H_2_O). All three of these crosslinks involve residues that are separated in C3 by distances (54, 46 and 44 Å) substantially greater than the theoretical cross-link limit (Fig. 5C). In spite of its movement towards MG7, ANA remains proximal to MG8 in C3(H_2_O), as evidenced by six MG8-ANA cross-links, four of which are C3(H_2_O)-unique (Fig. 5D). Furthermore, a C3(H_2_O)-unique cross-link between MG3 and ANA residues, 241^MG3^-670^ANA^, effectively replaces the C3-unique cross-link between these two domains, 267^MG3^-650^ANA^ (Fig. 5D, E). This observation suggests a new ANA-MG3 contact region has formed in C3(H_2_O).

Taken together, this cross-linking network in C3(H_2_O) between ANA on the one hand, and MG7, MG8 and MG3 on the other, places the ANA domain close to a hypothetical region of C3(H_2_O) where MG7, MG8 and MG3 converge, as they do in the C3b crystal structure (Fig. 5F, H). This new arrangement could be accomplished by migration of MG8 towards MG3 and a shift of the MG8-MG7 interface.

In the case of C3b formation, comparison of the crystal structures imply that the α’-NT segment relocates from its position in C3 to the other side of the molecule. Four cross-links from Ser727 in C3b supported this relocation. But none of these cross-links were observed in C3(H_2_O). Instead, the relative position of ANA and α-NT in C3(H_2_O) may be inferred to be similar to that in C3, on the basis of three C3/C3(H_2_O)-mutual cross-links between residues in ANA and α-NT (Fig. 5G). Presumably, the presence of the ANA domain in C3(H_2_O) prevents the relocation of the α’-NT segment to the opposite side of the molecule.

Moreover, the observation of C3-unique cross-link 882^MG7^-1539^C345C^ and C3b-unique cross-links 1479^Anchor^-1573^c3^4^5c^ and 1346^MG8^-1475^Anchor^ imply that in C3(H_2_O), the arrangement of the neck and the head (Fig. S1) is identical to that of neither C3 nor C3b.

In summary (Fig. 5I), inspection of QCLMS data for a three-way comparison allows us to conclude that C3(H_2_O) adopts a C3b-like conformation in terms of the relative arrangement of its TED and CUB domains. The presence of the ANA domain at the N terminus of the a-chain restricts the rearrangements possible in C3(H_2_O) thus resulting in a unique domain architecture within the α-chain that is different from the ones in C3 and C3b. We next sought to build a structural model of C3(H_2_O) based on the structures of C3 and C3b and the differences between them in terms of intramolecular cross-links.

### Integrative Modeling of C3(H_2_O)

In addition to structural inferences drawn from manual inspection and interpretation of crosslinks, we used IMP to compute the models from the data (46). Our QCLMS data included two key observations that allowed us to model the 3D structure of C3(H_2_O). First, the 48 cross-links that are mutual to C3, C3b and C3(H_2_O) confirmed that the structures of individual domains, together with the architecture of the β-chain, are largely preserved during the structural transitions that accompany C3 activation. Hence, we were able to treat individual domains, and the entire key-ring like core structure contributed by the β-chain (see Fig. 6A) as rigid bodies with known structures (from the crystal structure of C3) when modeling the structure of C3(H_2_O). Second, a total of 85 quantified cross-links (Supplemental Table S3) provided high-confidence distance restraints between pairs of residues in C3(H_2_O), which allowed for assembling individual rigid bodies into a 3D model of C3(H_2_O). Precise cross-linked sites were further assessed by manually inspecting the supporting fragmentation spectra, considering only residues K, S, T, Y and the protein N-termini as possible sites. If multiple possible sites were present in a peptide they were disambiguated by help of back-bone fragmentation events and in some cases using the chromatographic behavior of the cross-linked peptides as supporting information (examples shown as Supplemental Fig. S2 in Supplemental File). To assess our ability to build accurate 3D models in this way, we also built cross-link based models of C3 and C3b using the same approach and compared these against their crystal structures.

The models of C3, C3b, and C3(H_2_O) relied on the cross-link datasets, the crystallographic structures of domains in C3, the known disulfide bonds, the known primary structure, and the excluded volume, encoded in a Bayesian scoring function (*Experimental procedures*). The domains were represented by rigid-bodies connected by flexible linkers (Fig. 6A). We sampled the models by generating random translations and rotations of the rigid bodies, followed by the analysis of the best-scoring models (44–48). The resulting models of C3 and C3b (Fig. 6B, C) satisfied all of the input cross-link-derived restraints. Their architectures corresponded well to the respective crystal structures (Fig. 6E) as reflected in the accuracy (C-α r.m.s. deviation of the solutions with respect to the crystallographic structures) of approximately 11Å in both cases. In the case of C3(H_2_O), solutions with the best scores fall into two distinct clusters (A and B) corresponding to two alternative configurations of domains MG8 and ANA with respect to the other “shoulder” domains (Supplemental Fig. S3A, B). Cluster B better satisfies the cross-linking data (Supplemental Fig. S3C, D). The solutions in cluster B have a precision of 15 Å and satisfied 95% of the input cross-link-derived restraints. The four crosslinks that were not satisfied were 44^MG1^-1181^TED^, 44^MG1^-1195^TED^, 267^MG3^-1049^MG8^ and 727^α-NT^-1567^C345C^ (Fig. 6F). 44^MG1^-1181^TED^ and 44^MG1^-1195^TED^ could reflect the aforementioned mobility of TED with respect to the MG1-6 core. 267^MG3^-1049^MG8^ and 727^α-NT^-1567^C345C^ may reflect conformational changes within domains that were not allowed for in our rigid body-based modeling approach. For example, residue 1409^MG8^ lies within a pap-paa motif that exhibits different conformations between the structures of C3 (PDB|2A73) and C3b (PDB|2I07). The aNT segment (727-745) connecting the ANA and MG6 domains, appeared as a flexible loop in the C3 crystal structure yet had been treated as a rigid body, together with MG1-6 domains, for modeling purposes. It is also possible that the C345C domain is flexible relative to the shoulder region. Overall, the C3(H_2_O) models agree well with inferences based on manual inspection of cross-links (Fig. 5I) discussed in earlier.

## Discussion

The low-rate spontaneous activation of C3 occurs *via* a concerted process consisting of the hydrolysis of a thioester bond and a putative rearrangement of protein domains. This process is responsible for the ubiquitous and constitutive presence of C3(H_2_O) in plasma. It is the presence of C3(H_2_O) that maintains the alternative pathway of complement in “tick-over mode”, which is essential for the remarkable, near-instantaneous, response of complement to infection or danger first noted more than 100 years ago (56, 57). Unlike in the cases of C3 and C3b, no crystal structure of C3(H_2_O) has been reported and therefore the structural basis for spontaneous C3 activation is unproven.

In the current study we utilized QCLMS to reveal structural differences and similarities between C3(H_2_O), its progenitor C3, and its functional analogue C3b. This allowed us to infer the domain architecture of C3(H_2_O). We also combined QCMLS with structural knowledge, from crystallography, about the domains of C3 to build a computational model of the 3D structure of C3(H_2_O) with a precision of about 15 Å.

In QCMLS, pairs of amino acid residues within peptides that are tethered by a bifunctional cross-linker are identified using mass spectrometry. A potential source of error here is the possibility of misidentifying which residue within a peptide participates in the cross-link. A lead peptide-spectrum-match (PSM) is included in the supplement for every linked residue pair, showing no sign of abundant presence of this error. The impact of such errors on our findings is likely to be negligible for several reasons. First, our primary concern is modeling the spatial arrangement of protein domains (of between 70 and 300 residues) within a large (~180 kDa) multiple domain structure; hence misallocation, within a few residues, of a small number of cross-linked sites will not have significant consequences. Second, our findings are based on multiple, mutually corroborating cross-links. Third, a recent study using a similar approach reported that randomly altering cross-linked sites within an 11-residue window, had few repercussions (58). Moreover, to validate our procedure, we generated cross-link-derived models of C3 and C3b structures with an accuracy (compared to crystal structures) of about 11 Å. These results demonstrate the capacity of QCLMS to produce a set of self-consistent and highly informative distance restraints for proteins in solution.

We have thus provided experimental evidence that C3(H_2_O) adopts a C3b-like spatial arrangement of TED and CUB domains relative to the unchanging key ring-like core structure of MGs 1-6 (Fig. 4B). We further show that the rearrangement of the remaining, ANA, MG7, MG8, Anchor and C345C domains results in a unique conformation at the shoulder, neck and head regions of C3(H_2_O), which may be considered intermediate between those of C3 and C3b. We observed cross-links between TED and MG1 that are mutual to C3b and C3(H_2_O)b, but are not compatible with a single conformation of either protein. These cross-links imply that TED is mobile relative to the core of the protein. This observation agrees with several other strands of evidence for TED mobility in C3b (25, 54, 55) and shows there is a similar level of TED mobility in C3(H_2_O).

Our integrative model of C3(H_2_O) explains previously reported observations (8, 9, 21, 25, 27). A hydrogen/deuterium-exchange study reported that significant repositioning or reorientation of the CUB and TED domains accompanies the formation of C3(H_2_O) from C3. Images of C3(H_2_O) obtained by negative-stain transmission electron microscopy, show a predominance of C3b-like conformers in which TED had migrated from the shoulders of the molecule to its feet. Moreover, the similar structures of C3(H_2_O) and C3b explain their functional similarities. Both can bind factor H, and both act as a platform for binding of factor B and the subsequent factor D-mediated cleavage of factor B to Bb, although C3(H_2_O) is less effective in this role than C3b (see below) (59).

To explain the QCLMS data an adjustment is needed of a previous structural model of C3(H_2_O) based on EM images. In this model the ANA domain at the N terminus of the α-chain was proposed to migrate, together with the α’-NT-equivalent segment, from the MG8-MG3 side of the C3 structure to the opposite, MG7, side in C3(H_2_O) (25) (as does the nascent α’-NT of C3b). In contrast, our data indicated that the α’-NT-equivalent segment does not migrate across the structure to become exposed on the other side of C3(H_2_O). This implies that this segment of C3(H_2_O) would not contribute to the binding site for FH and FB as does α’-NT in C3b. These potential differences in the FH-binding and FB-binding surfaces (Fig. 5I) are consistent with the reportedly lower affinity of factor B for C3(H_2_O), compared to C3b (59), and may also pertain to the lower efficacy of factor H in deactivation of the C3(H_2_O)Bb complex (60). Taken together, our data placed the ANA domain at the contact point of the MG3, MG7 and MG8 domains (Fig. 5H, I, 6D), and on the opposite face of the molecule to the TED. Interestingly, the EM images (25) of C3(H_2_O) in complex with a FAb that binds to an epitope within the ANA domain, support the ANA location in our model.

It has been hypothesized that the ANA domain of C3 works as a molecular safety catch; its excision by proteolysis removes steric constraints on the domain rearrangements required for C3b formation (24). In agreement with this hypothesis, our data show that in C3(H_2_O) the ANA domain is displaced (Fig. 5H, I, 6D), and this has a similar effect to excision on the surrounding domains. Interestingly, an inactive C3(H_2_O)/C3 intermediate has been reported, with a hydrolyzed thioester, which resembles native C3 in overall shape (20, 25). Taken together these observations imply that it is the displacement of ANA that serves as the trigger for the dramatic structural rearrangements that accompany formation of active C3(H_2_O).

Our analysis of C3(H_2_O) has demonstrated that a hitherto unknown structure can be determined by QCLMS data and modeling, widening the application range of this technology.

## Acknowledgments

We thank Christoph Schmidt, Carla Clark and Paul McLaughlin for helpful discussions. We thank Sven Giese for helping with annotated MS2 spectra. We acknowledge the PRIDE team for the deposition of our data to the ProteomeXchange Consortium. We thank Tru Huynh for help with the computer cluster at Institut Pasteur, and Ben Webb for support with the Integrative Modeling Platform. This work was supported by the Wellcome Trust (PNB: 081179, JR: 103139, Centre core grant: 092076, instrument grant: 108504), BBSRC (BB/I007946/1), the NIH R01 GM083960 and P41 GM083960 grants to AS, and the European Research Council (MN: FP7-IDEAS-ERC 294809). JR is a Senior Research Fellow of the Wellcome Trust.

## References

1. Ricklin, D., Hajishengallis, G., Yang, K., and Lambris, J. D. (2010) Complement: a key system for immune surveillance and homeostasis. Nat Immunol 11, 785–797

2. Walport, M. J. (2001) Complement. First of two parts. N Engl J Med 344, 1058–1066

3. Law, S. K., Lichtenberg, N. A., and Levine, R. P. (1979) Evidence for an ester linkage between the labile binding site of C3b and receptive surfaces. J Immunol 123, 1388–1394

4. Tack, B. F., Harrison, R. A., Janatova, J., Thomas, M. L., and Prahl, J. W. (1980) Evidence for presence of an internal thiolester bond in third component of human complement. Proc Natl Acad Sci U S A 77, 5764–5768

5. Isaac, L., Aivazian, D., Taniguchi-Sidle, A., Ebanks, R. O., Farah, C. S., Florido, M. P., Pangburn, M. K., and Isenman, D. E. (1998) Native conformations of human complement components C3 and C4 show different dependencies on thioester formation. Biochem J 329 (Pt 3), 705–712

6. Pangburn, M. K., and Muller-Eberhard, H. J. (1980) Relation of putative thioester bond in C3 to activation of the alternative pathway and the binding of C3b to biological targets of complement. J Exp Med 152, 1102–1114

7. Gros, P., Milder, F. J., and Janssen, B. J. (2008) Complement driven by conformational changes. Nat Rev Immunol 8, 48–58

8. Isenman, D. E., Kells, D. I., Cooper, N. R., Muller-Eberhard, H. J., and Pangburn, M. K. (1981) Nucleophilic modification of human complement protein C3: correlation of conformational changes with acquisition of C3b-like functional properties. Biochemistry 20, 4458–4467

9. Li, K., Gor, J., and Perkins, S. J. (2010) Self-association and domain rearrangements between complement C3 and C3u provide insight into the activation mechanism of C3. Biochem J 431, 63–72

10. Pangburn, M. K., Schreiber, R. D., and Muller-Eberhard, H. J. (1981) Formation of the initial C3 convertase of the alternative complement pathway. Acquisition of C3b-like activities by spontaneous hydrolysis of the putative thioester in native C3. J Exp Med 154, 856–867

11. Janssen, B. J., Christodoulidou, A., McCarthy, A., Lambris, J. D., and Gros, P. (2006) Structure of C3b reveals conformational changes that underlie complement activity. Nature 444, 213–216

12. Alsenz, J., Becherer, J. D., Nilsson, B., and Lambris, J. D. (1990) Structural and functional analysis of C3 using monoclonal antibodies. Curr Top Microbiol Immunol 153, 235–248

13. Becherer, J. D., Alsenz, J., and Lambris, J. D. (1990) Molecular aspects of C3 interactions and structural/functional analysis of C3 from different species. Curr Top Microbiol Immunol 153, 45–72

14. Becherer, J. D., and Lambris, J. D. (1988) Identification of the C3b receptor-binding domain in third component of complement. J Biol Chem 263, 14586–14591

15. Forneris, F., Ricklin, D., Wu, J., Tzekou, A., Wallace, R. S., Lambris, J. D., and Gros, P. (2010) Structures of C3b in complex with factors B and D give insight into complement convertase formation. Science 330, 1816–1820

16. Lambris, J. D., Avila, D., Becherer, J. D., and Muller-Eberhard, H. J. (1988) A discontinuous factor H binding site in the third component of complement as delineated by synthetic peptides. J Biol Chem 263, 12147–12150

17. Lambris, J. D., Lao, Z., Oglesby, T. J., Atkinson, J. P., Hack, C. E., and Becherer, J. D. (1996) Dissection of CR1, factor H, membrane cofactor protein, and factor B binding and functional sites in the third complement component. J Immunol 156, 4821–4832

18. Morgan, H. P., Schmidt, C. Q., Guariento, M., Blaum, B. S., Gillespie, D., Herbert, A. P., Kavanagh, D., Mertens, H. D., Svergun, D. I., Johansson, C. M., Uhrin, D., Barlow, P. N., and Hannan, J. P. (2011) Structural basis for engagement by complement factor H of C3b on a self surface. Nat Struct Mol Biol 18, 463–470

19. Ricklin, D., and Lambris, J. D. (2007) Complement-targeted therapeutics. Nat Biotechnol 25, 1265–1275

20. Pangburn, M. K. (1992) Spontaneous reformation of the intramolecular thioester in complement protein C3 and low temperature capture of a conformational intermediate capable of reformation. J Biol Chem 267, 8584–8590

21. Hack, C. E., Paardekooper, J., Smeenk, R. J., Abbink, J., Eerenberg, A. J., and Nuijens, J. H. (1988) Disruption of the internal thioester bond in the third component of complement (C3) results in the exposure of neodeterminants also present on activation products of C3. An analysis with monoclonal antibodies. J Immunol 141, 1602–1609

22. Isenman, D. E. (1983) Conformational changes accompanying proteolytic cleavage of human complement protein C3b by the regulatory enzyme factor I and its cofactor H. Spectroscopic and enzymological studies. J Biol Chem 258, 4238–4244

23. Isenman, D. E., and Cooper, N. R. (1981) The structure and function of the third component of human complement-I. The nature and extent of conformational changes accompanying C3 activation. Mol Immunol 18, 331–339

24. Janssen, B. J., Huizinga, E. G., Raaijmakers, H. C., Roos, A., Daha, M. R., Nilsson-Ekdahl, K., Nilsson, B., and Gros, P. (2005) Structures of complement component C3 provide insights into the function and evolution of immunity. Nature 437, 505–511

25. Nishida, N., Walz, T., and Springer, T. A. (2006) Structural transitions of complement component C3 and its activation products. Proc Natl Acad Sci U S A 103, 19737–19742

26. Perkins, S. J., and Sim, R. B. (1986) Molecular modelling of human complement component C3 and its fragments by solution scattering. Eur J Biochem 157, 155–168

27. Winters, M. S., Spellman, D. S., and Lambris, J. D. (2005) Solvent accessibility of native and hydrolyzed human complement protein 3 analyzed by hydrogen/deuterium exchange and mass spectrometry. J Immunol 174, 3469–3474

28. Fischer, L., Chen, Z. A., and Rappsilber, J. (2013) Quantitative cross-linking/mass spectrometry using isotope-labelled cross-linkers. Journal of proteomics 88, 120–128

29. Schmidt, C., Zhou, M., Marriott, H., Morgner, N., Politis, A., and Robinson, C. V. (2013) Comparative cross-linking and mass spectrometry of an intact F-type ATPase suggest a role for phosphorylation. Nature communications 4, 1985

30. Kukacka, Z., Rosulek, M., Strohalm, M., Kavan, D., and Novak, P. (2015) Mapping protein structural changes by quantitative cross-linking. Methods 89, 112–120

31. Walzthoeni, T., Joachimiak, L. A., Rosenberger, G., Rost, H. L., Malmstrom, L., Leitner, A., Frydman, J., and Aebersold, R. (2015) xTract: software for characterizing conformational changes of protein complexes by quantitative cross-linking mass spectrometry. Nat Methods 12, 1185–1190

32. Chen, Z. A., Fischer, L., Tahir, S., Bukowski-Wills, J.-C., Barlow, P. N., and Rappsilber, J. (2016) Quantitative cross-linking/mass spectrometry reveals subtle protein conformational changes. bioRxiv doi.10.1101/055418

33. Tomko, R. J. Jr.,, Taylor, D. W., Chen, Z. A., Wang, H. W., Rappsilber, J., and Hochstrasser, M. (2015) A Single alpha Helix Drives Extensive Remodeling of the Proteasome Lid and Completion of Regulatory Particle Assembly. Cell 163, 432–444

34. Sanchez-Corral, P., Anton, L. C., Alcolea, J. M., Marques, G., Sanchez, A., and Vivanco, F. (1989) Separation of active and inactive forms of the third component of human complement, C3, by fast protein liquid chromatography (FPLC). J Immunol Methods 122, 105–113

35. Maiolica, A., Cittaro, D., Borsotti, D., Sennels, L., Ciferri, C., Tarricone, C., Musacchio, A., and Rappsilber, J. (2007) Structural analysis of multiprotein complexes by cross-linking, mass spectrometry, and database searching. Mol Cell Proteomics 6, 2200–2211

36. Ishihama, Y., Rappsilber, J., and Mann, M. (2006) Modular stop and go extraction tips with stacked disks for parallel and multidimensional Peptide fractionation in proteomics. J Proteome Res 5, 988–994

37. Rappsilber, J., Mann, M., and Ishihama, Y. (2007) Protocol for micro-purification, enrichment, pre-fractionation and storage of peptides for proteomics using StageTips. Nat Protoc 2, 1896–1906

38. Rappsilber, J., Ishihama, Y., and Mann, M. (2003) Stop and go extraction tips for matrix-assisted laser desorption/ionization, nanoelectrospray, and LC/MS sample pretreatment in proteomics. Anal Chem 75, 663–670

39. Ishihama, Y., Rappsilber, J., Andersen, J. S., and Mann, M. (2002) Microcolumns with self-assembled particle frits for proteomics. J Chromatogr A 979, 233–239

40. Cox, J., and Mann, M. (2008) MaxQuant enables high peptide identification rates, individualized p.p.b.-range mass accuracies and proteome-wide protein quantification. Nat Biotechnol 26, 1367–1372

41. Thakur, S. S., Geiger, T., Chatterjee, B., Bandilla, P., Frohlich, F., Cox, J., and Mann, M. (2011) Deep and highly sensitive proteome coverage by LC-MS/MS without prefractionation. Mol Cell Proteomics 10, M110 003699

42. DeLano, W. L. (2002) The PyMOL Molecular Graphics System. DeLano Scientific, San Carlos, CA, USA

43. Alber, F., Dokudovskaya, S., Veenhoff, L. M., Zhang, W., Kipper, J., Devos, D., Suprapto, A., Karni-Schmidt, O., Williams, R., Chait, B. T., Rout, M. P., and Sali, A. (2007) Determining the architectures of macromolecular assemblies. Nature 450, 683–694

44. Erzberger, J. P., Stengel, F., Pellarin, R., Zhang, S., Schaefer, T., Aylett, C. H., Cimermancic, P., Boehringer, D., Sali, A., Aebersold, R., and Ban, N. (2014) Molecular architecture of the 40SeIF1eIF3 translation initiation complex. Cell 158, 1123–1135

45. Robinson, P. J., Trnka, M. J., Pellarin, R., Greenberg, C. H., Bushnell, D. A., Davis, R., Burlingame, A. L., Sali, A., and Kornberg, R. D. (2015) Molecular architecture of the yeast Mediator complex. eLife 4

46. Russel, D., Lasker, K., Webb, B., Velazquez-Muriel, J., Tjioe, E., Schneidman-Duhovny, D., Peterson, B., and Sali, A. (2012) Putting the pieces together: integrative modeling platform software for structure determination of macromolecular assemblies. PLoS biology 10, e1001244

47. Shi, Y., Fernandez-Martinez, J., Tjioe, E., Pellarin, R., Kim, S. J., Williams, R., Schneidman-Duhovny, D., Sali, A., Rout, M. P., and Chait, B. T. (2014) Structural characterization by cross-linking reveals the detailed architecture of a coatomer-related heptameric module from the nuclear pore complex. Mol Cell Proteomics 13, 2927–2943

48. Shi, Y., Pellarin, R., Fridy, P. C., Fernandez-Martinez, J., Thompson, M. K., Li, Y., Wang, Q. J., Sali, A., Rout, M. P., and Chait, B. T. (2015) A strategy for dissecting the architectures of native macromolecular assemblies. Nat Methods 12, 1135–1138

49. Molnar, K. S., Bonomi, M., Pellarin, R., Clinthorne, G. D., Gonzalez, G., Goldberg, S. D., Goulian, M., Sali, A., and DeGrado, W. F. (2014) Cys-scanning disulfide crosslinking and bayesian modeling probe the transmembrane signaling mechanism of the histidine kinase, PhoQ. Structure 22, 1239–1251

50. Rieping, W., Habeck, M., and Nilges, M. (2005) Inferential structure determination. Science 309, 303–306

51. Fernandez-Martinez, J., Phillips, J., Sekedat, M. D., Diaz-Avalos, R., Velazquez-Muriel, J., Franke, J. D., Williams, R., Stokes, D. L., Chait, B. T., Sali, A., and Rout, M. P. (2012) Structure-function mapping of a heptameric module in the nuclear pore complex. J Cell Biol 196, 419–434

52. Vizcaino, J. A., Deutsch, E. W., Wang, R., Csordas, A., Reisinger, F., Rios, D., Dianes, J. A., Sun, Z., Farrah, T., Bandeira, N., Binz, P. A., Xenarios, I., Eisenacher, M., Mayer, G., Gatto, L., Campos, A., Chalkley, R. J., Kraus, H. J., Albar, J. P., Martinez-Bartolome, S., Apweiler, R., Omenn, G. S., Martens, L., Jones, A. R., and Hermjakob, H. (2014) ProteomeXchange provides globally coordinated proteomics data submission and dissemination. Nat Biotechnol 32, 223–226

53. Pangburn, M. K., Schreiber, R. D., and Muller-Eberhard, H. J. (1977) Human complement C3b inactivator: isolation, characterization, and demonstration of an absolute requirement for the serum protein beta1 H for cleavage of C3b and C4b in solution. J Exp Med 146, 257–270

54. Alcorlo, M., Tortajada, A., Rodriguez de Cordoba, S., and Llorca, O. (2013) Structural basis for the stabilization of the complement alternative pathway C3 convertase by properdin. Proc Natl Acad Sci U S A 110, 13504–13509

55. Pechtl, I. C., Neely, R. K., Dryden, D. T., Jones, A. C., and Barlow, P. N. (2011) Use of time-resolved FRET to validate crystal structure of complement regulatory complex between C3b and factor H (N terminus). Protein Sci 20, 2102–2112

56. Ehrlich, P. (1899) Zur Theorie der Lysin Wirkung. Berl. Klin. Wochenschr, 6–9

57. Metschnikow, I. I. (1902) Immunität bei Infektionskrankheiten, Fischer, Frankfurt

58. Belsom, A., Schneider, M., Fischer, L., Brock, O., and Rappsilber, J. (2016) Serum Albumin Domain Structures in Human Blood Serum by Mass Spectrometry and Computational Biology. Mol Cell Proteomics 15, 1105–1116

59. Pryzdial, E. L., and Isenman, D. E. (1988) A thermodynamic study of the interaction between human complement components C3b or C3(H_2_O) and factor B in solution. J Biol Chem 263, 1733–1738

60. Bexborn, F., Andersson, P. O., Chen, H., Nilsson, B., and Ekdahl, K. N. (2008) The tick-over theory revisited: formation and regulation of the soluble alternative complement C3 convertase (C3(H_2_O)Bb). Mol Immunol 45, 2370–2379

